# Designed Soluble Notch Agonist Drives Human Ameloblast Maturation for Tooth Regeneration

**DOI:** 10.1101/2025.04.03.646929

**Authors:** Anjali P. Patni, Rubul Mout, Rachel Moore, Ammar Alghadeer, George Q Daley, David Baker, Julie Mathieu, Hannele Ruohola-Baker

**Affiliations:** Department of Oral Health Sciences, University of Washington, School of Dentistry, Seattle, WA 98109, USA; Department of Biochemistry, University of Washington, School of Medicine, Seattle, WA 98195, USA; Institute for Stem Cell and Regenerative Medicine, University of Washington, School of Medicine, Seattle, WA 98109, USA; Stem Cell Program, Boston Children’s Hospital, Boston, MA 02115, USA; Department of Biological Chemistry and Molecular Pharmacology, Blavatnik Institute Harvard Medical School, Boston, MA 02115, USA; Harvard Stem Cell Institute, Harvard University, Cambridge, MA 02138, USA; Department of Comparative Medicine, University of Washington, School of Medicine, Seattle, WA 98195, USA; Department of Biomedical Dental Sciences, Imam Abdulrahman Bin Faisal University, College of Dentistry, Dammam 31441, Saudi Arabia; Institute for Protein Design, University of Washington, Seattle, WA 98195, USA; Howard Hughes Medical Institute, University of Washington, Seattle, WA 98195, USA

## Abstract

Enamel, the hardest material in the human body, is required to protect our living organ, tooth. However, over 90% of adults have lost or damaged enamel and cannot regenerate the protective structure due to lack of enamel producing cells, ameloblasts. iPSC derived mature Ameloblasts (iAM) have promise in future regenerative dentistry. Today it is not known why iAM maturation requires intimate contact with the dentin producing cell type, odontoblast. Here we reveal that one of the critical signaling ligands emanating from odontoblasts for ameloblast maturation is Delta, the ligand for Notch receptor. We showed that our designed, soluble Notch agonist can induce iAM organoid maturation in an unprecedented manner, without interactions with odontoblast layer. This novel maturation procedure enables us to analyze the specific requirements of DLX3 function in ameloblasts, independent of its known function in odontoblasts. We now show that DLX3, the gene associated with Amelogenesis Imperfecta, is required on a cell-autonomous manner in ameloblasts for the expression of Enamelin and MMP20.

## Introduction

From the teeth of Krakens, giant squids in deep sea to elephant’s ivory tusks in savanna, these living ectodermal structures are indispensable organs for animal survival. In mammals and reptiles alike, teeth are protected by an outer calcified layer called enamel. While some animals can regenerate continuously this protective enamel layer, humans cannot. In human developing tooth the enamel secreting cells, ameloblasts undergo apoptosis at the time of tooth eruption resulting in lost regenerative capacity of the protective layer in our aging body.

Amelogenesis Imperfecta (AI) is a genetic disorder that causes impaired enamel development due to defects in the specialized cells secreting the protective enamel layer, resulting in fragile teeth with irregular shape, thickness, and pigmentation^1^. Today we still do not have the capacity to generate the mature secretory cell type, Ameloblast, hampering regenerative restoration or other therapeutic interventions.

Recent progress in regenerative dentistry has highlighted the application of human induced pluripotent stem cells (hiPSCs) to generate induced ameloblast cells (iAMs). These iAMs possess significant potential for upcoming approaches aimed at enamel repair and regeneration. If iAMs can be differentiated into functional ameloblasts, they could serve as a viable source of cells for enamel regeneration. However, today the maturation of iAMs remains a significant hurdle, primarily due to a necessity for understanding the specific mechanisms and signals required for their development. One particularly intriguing element is the crucial requirement for close interaction of AMs with odontoblasts (OBs), the dentin-producing cells^2^. Odontoblast–ameloblast crosstalk is supported by studies of SP7 (also known as Osterix), a transcription factor specifically expressed in odontoblasts and osteoblasts but not in ameloblasts^3^. Conditional knockout mouse models targeting SP7-expressing cells (e.g., *Osx-Cre; Macf1^fl/fl^*, *Osx-Cre; Stat3^fl/fl^*, *Osx-Cre; Cbf*β*^fl/fl^*) ^4, 5, 6–11^ frequently exhibit both dentin and enamel defects, suggesting that compromised SP7-expressing odontoblasts may indirectly affect ameloblast activity and pointing to potential crosstalk between these two cell populations.

One critical cell-cell communication signaling pathway, Notch pathway is a highly conserved cell signaling mechanism known to govern multiple stages in cellular differentiation^12,13^. As an example, it plays a pivotal role in maintaining the balance between progenitor cell proliferation and differentiation, ensuring proper tissue development and homeostasis^12^. The Notch pathway operates through direct cell-cell communication, wherein Notch receptors (Notch1–4) on a responding cell interact with ligands (Jagged1/2 and Delta-like ligands DLL1, DLL3, and DLL4) presented on the surface of neighboring cells. Upon ligand-receptor engagement, the Notch receptor undergoes two sequential proteolytic cleavages. The first cleavage, mediated by ADAM-family metalloproteases, releases the extracellular domain of the receptor, while the second cleavage, facilitated by the γ-secretase complex, liberates the Notch intracellular domain (NICD). Once freed, the NICD translocate to the nucleus, forming a complex with the transcriptional regulators RBP-J/γ-CSL and MAML. These interactions drive the transcription of Notch target genes, including members of the Hes and Hey families, which play critical roles in cell differentiation, proliferation, and survival. This tightly regulated cascade ensures that Notch signaling is activated in response to direct cell-cell interaction, enabling specific and localized effects on tissue morphogenesis. Interestingly, recent genome wide association studies (GWAS) have identified a significant link between a single nucleotide polymorphism (SNP) within an intron of ADAMTS9 gene and amelogenesis imperfecta^14,15.^

We have previously used single-cell sequencing of human fetal ameloblasts (AM) to develop a human iPSC-based induced AM (iAM) maturation protocol^16^ to generate induced secretory ameloblasts (isAM). In this protocol, Ameloblasts require interactions with odontoblasts (co-culture) to exhibit mineralization and expression of Amelogenin (AMELX), and Enamelin (ENAM)^2^. In the present study we identify the critical signal emanating from OB that activates Notch signaling resulting in maturation of ameloblasts. We showed that our computer-designed soluble Notch activator^17^ can generate mature ameloblasts in the absence of odontoblasts, enabling us to identify the mode of function of the amelogenesis imperfecta gene, DLX3.

## Results

### Notch pathway during Ameloblast maturation

iPSC-derived iAM requires a co-culture with iOB for the maturation into secretory ameloblasts ^2^. To identify the critical signaling pathways emanating from odontoblasts for AM maturation we performed TopPath algorithm analysis for previous single-cell-sequencing data from human fetal tissues^2,18^. This analysis revealed Notch as a candidate signaling pathway involved in the crosstalk between OB and AM during the AM maturation stage (Fig.1A-B). While other pathways (FGF, WNT and EGF) were also identified as potentially important signaling pathways at this stage, their contribution from OB to sAM signaling was not predicted by the talklr R package^19^ to be as prominent as Notch pathway (Figure 1A-B; Figure S1B-D).

**Figure 1.**
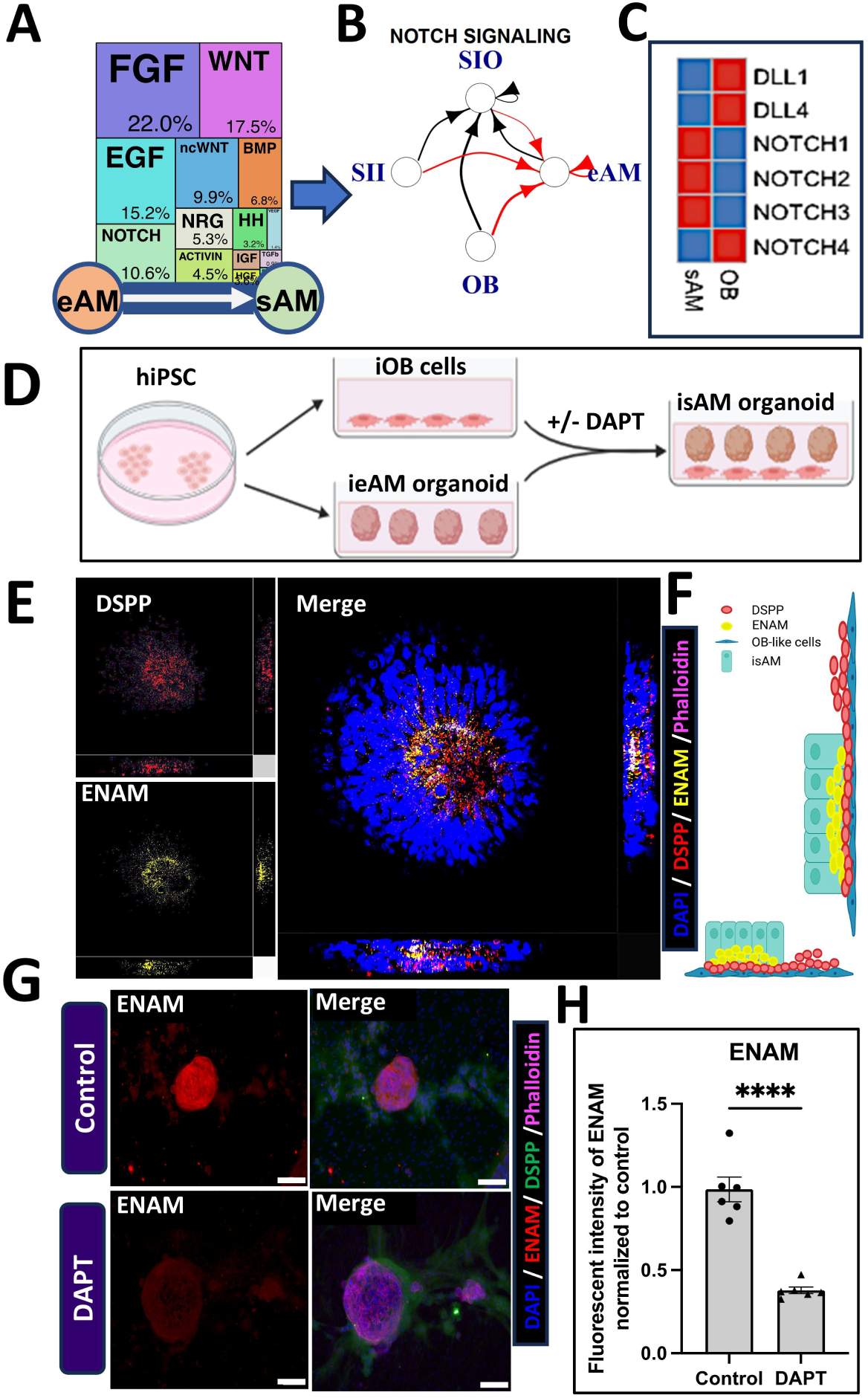
Notch pathway activation is required for ameloblast maturation in co-culture with odontoblasts. (A) TopPath pathway analysis of single cell sequencing data from fetal tissues, identifying key signaling pathways involved in odontoblast (OB) to ameloblast (AM) communication. The Notch pathway (10.6%) is highlighted as a major contributor to AM maturation, alongside other pathways such as FGF, WNT, EGF, and BMP. (B) Schematic model depicting Notch signaling interactions between OBs and AMs, showing ligand-receptor interactions focusing on incoming signal to eAM (red arrows). The width of the arrows is proportional to the number of interactions. Black arrow indicates other possible interaction not considered in the analysis of TopPath. (C) Heatmap representation of Notch ligands (DLL1, DLL4) and Notch receptors (NOTCH1, NOTCH2, NOTCH3, NOTCH4) obtained from the single-cell data at the late timepoints during human fetal development (20-22gw). (D) Differentiation and co-culture model for generating induced ameloblasts (iAMs) from human iPSCs. Induced odontoblasts (iOBs) are co-cultured with induced early ameloblast (ieAM) organoids to promote their maturation into secretory ameloblasts (isAMs), with and without DAPT-mediated Notch inhibition. (E) 3D confocal images of co-cultured iAM organoids showing ENAM (yellow) and DSPP (red) expression, confirming ameloblast maturation in co-culture. Merged images include DAPI (nuclei, blue) and Phalloidin (cytoskeleton, purple). (F) Schematic representation summarizing the observed expression patterns of DSPP (red) and ENAM (yellow) in differentiated OB-like and AM-like cells within the co-culture. (G) Comparative immunofluorescence staining of ENAM in Control (top) vs. DAPT-treated (bottom) conditions, revealing reduced ENAM expression upon Notch inhibition. (H) Quantification of ENAM fluorescence intensity, normalized to control, showing a significant reduction in ENAM expression in the DAPT-treated group (****p < 0.0001).

To evaluate the expression of Notch ligands and receptors in OB and sAM, we analyzed the gene expression levels of Notch and Delta in these cell types. We generated a heatmap by averaging the expression levels of OB and AM clusters from the single-cell data at the late timepoints during human fetal development (20-22gw)^2^ (File S1). We found that the ligands (DLL1 and DLL4) were predominantly expressed in the OB, while the Notch receptors (NOTCH1, NOTCH2, and NOTCH3) were primarily present on the sAM side (Figure 1C), suggesting that Delta from Odontoblasts might activate Notch in Ameloblasts.

### Notch inhibitor affects enamelin expression in AM and OB co-culture system

To study the interaction between OB and AM we optimized our previously published co-culture protocol by developing a serum free protocol for iOB generation^18^ (Figure S1A). This process is enhanced by activating FGF signaling through a novel C6-mb7 FGFR1/2c agonist^20^ and Hedgehog signaling via SAG, supporting their maturation into functional iOBs by day 25 (Figure S1A). These mature iOBs are plated as a monolayer in Matrigel to create a physiological substrate for downstream experiments. To generate the co-culture system, we differentiated hiPSC derived ameloblast (iAM) organoids as done before^2^. Ameloblast differentiation is initiated by inducing oral epithelial lineage from hiPSCs under defined serum-free conditions from day 0 to day 10. At day 16 the early ameloblasts (ieAM) were directed to three-dimensional (3D) organoids using low-adhesion plates and at day 24 the organoids are co-cultured with iOB monolayers in Matrigel, recreating the *in vivo* tooth development microenvironment (Figure 1D; Figure S1A). This co-culture system enables reciprocal interactions between iOB and ieAM, driving the latter’s progression to secretory ameloblast stage (isAM). This state is characterized by polarized cells secreting enamelin (ENAM), an essential protein for enamel formation, with polarity reversal of ieAM toward the iOB layer, indicative of interaction-driven maturation (Figure 1E-1F). We utilized the hiPSCs based ameloblast (iAM) and odontoblast differentiation (iOB) methods in a defined serum-free media approach to dissect Notch function in OB to AM signaling. To assess if Notch signaling is essential for ameloblast (AM) maturation, we used DAPT (N-[N-(3, 5-difluorophenacetyl)-l-alanyl]-s-phenylglycinet-butyl ester)^21^, a well-characterized small molecule inhibitor of γ-secretase. γ-secretase, with Presenilin as its catalytic subunit, is responsible for cleaving the Notch receptor. This cleavage releases the Notch intracellular domain (NICD), allowing it to translocate to the nucleus and activate target gene transcription involved in cellular differentiation. By inhibiting γ-secretase, DAPT blocks the release of NICD, disrupting Notch signaling. Prior studies have shown that Notch signaling is essential for the survival and maintenance of epithelial stem cells in the continuously growing mouse incisor^22^. Consistent with these findings, while iOB-iAM interaction drove ENAM secretion in the co-culture system, Notch inhibition via DAPT significantly reduced ENAM induction in this system (Figure 1G-1H), suggesting that active Notch signaling is crucial for AM maturation and enamelin protein synthesis in this model.

### Soluble Notch agonist

Notch activity has been implicated in the differentiation of ameloblast support cells in dental tissue development^23,24^. Specifically, Notch signaling mediates the divergence of the stratum intermedium lineage from the ameloblast lineage, highlighting its crucial role in orchestrating cell fate decisions in the enamel organ. To specifically dissect the function of Notch activity in ameloblast (AM) maturation, it is essential to achieve precise spatial and temporal control over Notch activation. In response to this challenge, we have utilized our soluble, computationally designed Notch activator^17^. This scaffold allows precise temporal regulation of Notch signaling, facilitating a detailed investigation into its role during AM maturation and enamel formation. The Notch activator scaffold (C3-DLL4) is a computationally designed^25 26,27^, multivalent, soluble protein complex capable of activating the Notch pathway (Figure 2A)^17^. This synthetic system features an engineered oligomeric structure^28–30^ composed of three repeating subunits conjugated to the Delta-like 4 extracellular domain (DLL4). By arranging the DLL4 ectodomain in a trimeric formation on a helical bundle, soluble C3-DLL4 mimics the natural ligand presentation required for Notch activation^17^. Specifically, C3-DLL4 configuration promotes interaction between neighboring cell surfaces, providing the necessary tethering and mechanical tension to activate the Notch pathway. Importantly, C3-DLL4 overcomes the limitations of generating mechanical tension by traditional immobilization methods, enabling effective Notch signaling by soluble ligand.

**Figure 2.**
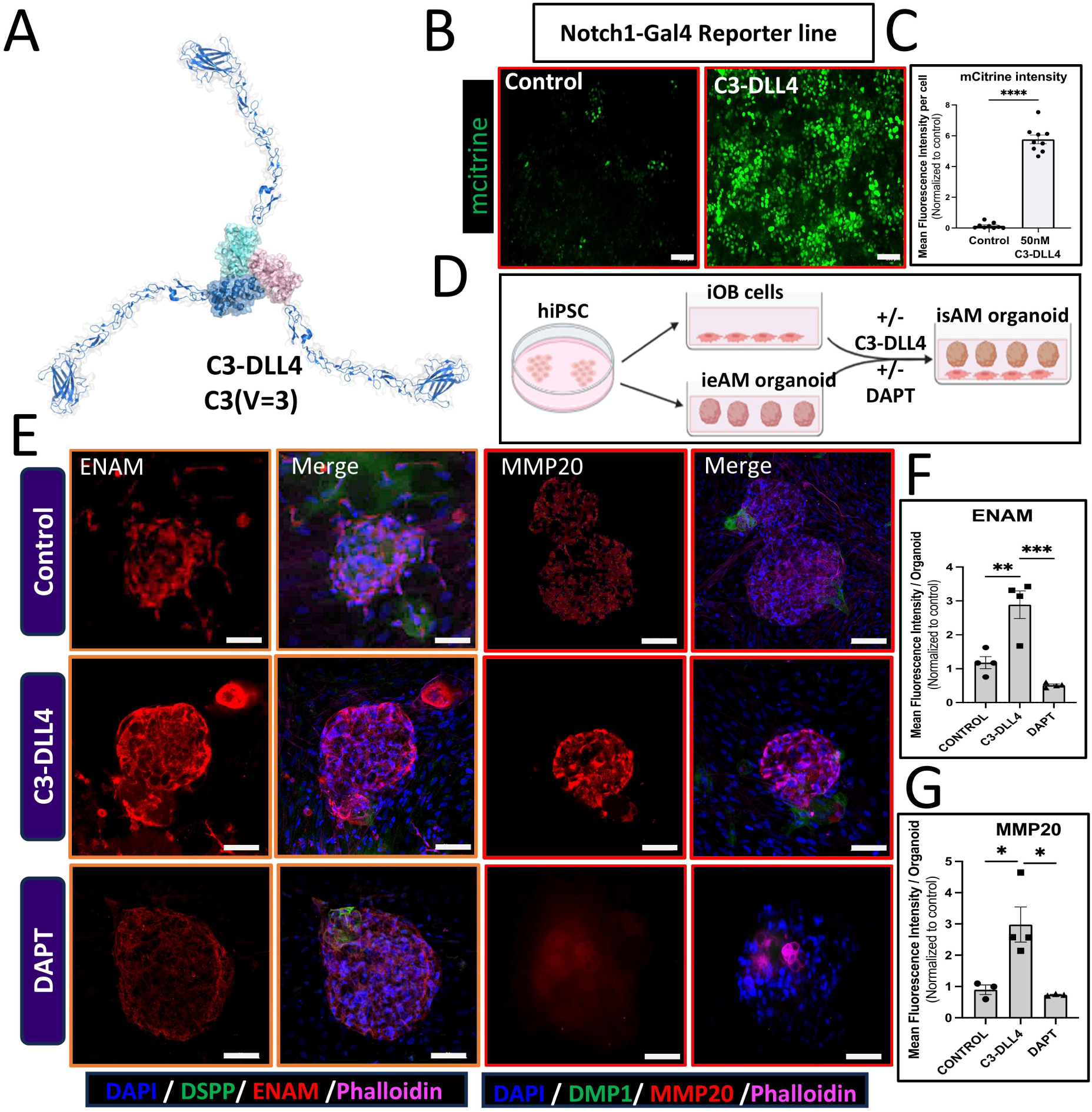
AI-designed Notch activator C3-DLL4 enhances ameloblast maturation in co-culture. (A) Structural model of the computationally designed C3-DLL4 protein complex, showing its trivalent configuration (C3(V=3)), which clusters Notch receptors to simulate ligand-receptor interactions in cell communication. (B) Notch1-Gal4 reporter assay in U2OS cells treated with control or C3-DLL4, showing nuclear mCitrine expression exclusively in C3-DLL4-treated cells, confirming Notch pathway activation. (C) Quantification of mCitrine fluorescence intensity per cell, revealing a significant increase in Notch activation upon C3-DLL4 treatment (****p < 0.0001) compared to control. (D) Schematic of the experimental setup, illustrating the differentiation of iOBs and iAMs, followed by co-culture with or without C3-DLL4 and DAPT treatment to assess Notch activation in ameloblast maturation. (E) Immunofluorescence staining of 3D iAM organoids treated with Control (top), C3-DLL4 (middle), and DAPT (bottom), showing ENAM (red) and MMP20 (red) expression. Control organoids exhibit moderate ENAM and MMP20 levels, while C3-DLL4-treated organoids show increased expression, and DAPT-treated organoids exhibit strongly reduced expression. Merged images include DAPI (nuclei, blue), DSPP/DMP1 (odontoblast markers, green), ENAM (red), and Phalloidin (cytoskeleton, purple). (F) Quantification of ENAM fluorescence intensity, normalized to control, demonstrating a significant increase in ENAM expression in C3-DLL4-treated organoids (\*\*\**p < 0.001*) and a significant reduction in DAPT-treated organoids (*p < 0.01). (G) Quantification of MMP20 fluorescence intensity, showing a significant increase in C3-DLL4-treated organoids (*p < 0.05) and a marked reduction in DAPT-treated organoids (*p < 0.05).

To evaluate the effectiveness of the conjugated C3-DLL4 in activating Notch signaling, we used the established Notch1-Gal4 reporter system^31^, which minimizes interference from endogenous Notch signals by replacing the NOTCH1 ankyrin repeat domain with Gal4 transcription activator. Upon binding to Delta, Notch receptor proteolytic processing releases a chimeric NICD1-Gal4 protein that can activate the engineered UAS-mCitrine gene. We showed that C3-DLL4 led to significant nuclear mCitrine expression, confirming Notch activation (Figure 2B-2C). Control cells treated with doxycycline did not display nuclear mCitrine, validating the specificity of C3-DLL4 in driving Notch activation. This demonstrated the effectiveness of soluble, computer designed C3-DLL4 to activate Notch receptor.

### Notch activation accelerates isAM maturation in co-culture system

We applied C3-DLL4 in our co-culture system, to assess if Notch activation could accelerate induced ameloblast maturation to isAM stage. We differentiated iOB and iAM independently, combined the cultures and treated the co-cultured cells with C3-DLL4 (50nM for 24 hours) (Figure 2D-E). Treatment with C3-DLL4 enhanced the expression of mature enamel matrix protein (EMP) markers, including ENAM and MMP20, indicating that active Notch signaling promotes isAM maturation (Figure 2E-G). These findings suggest that C3-DLL4-mediated Notch activation effectively enhances AM maturation in the co-culture model.

### C3-DLL4 accelerates iAM maturation without co-culture

We further investigated if C3-DLL4 could accelerate iAM maturation without close proximity to iOB (Figure 2A; Figure S2A-S2B). Given that our initial results identified Notch as a candidate mediator of ameloblast maturation, particularly driven by interactions with iOB-derived Delta ligands, we hypothesized that direct activation of Notch in iAM might bypass the need for OB co-culture conditions. To assess this hypothesis, we generated early ameloblasts (ieAM) in 3D suspension cultures enabling self-organization into ieAM organoids (Figure 3A; Figure S2B). This 3D environment supports early ameloblast maturation, mimicking the spatial context of natural tooth development. We applied C3-DLL4 to ieAM organoids (50nM, 24hours) and the organoids were maintained for an additional 7 days to evaluate long-term effects on maturation. (Figure 3A).

**Figure 3.**
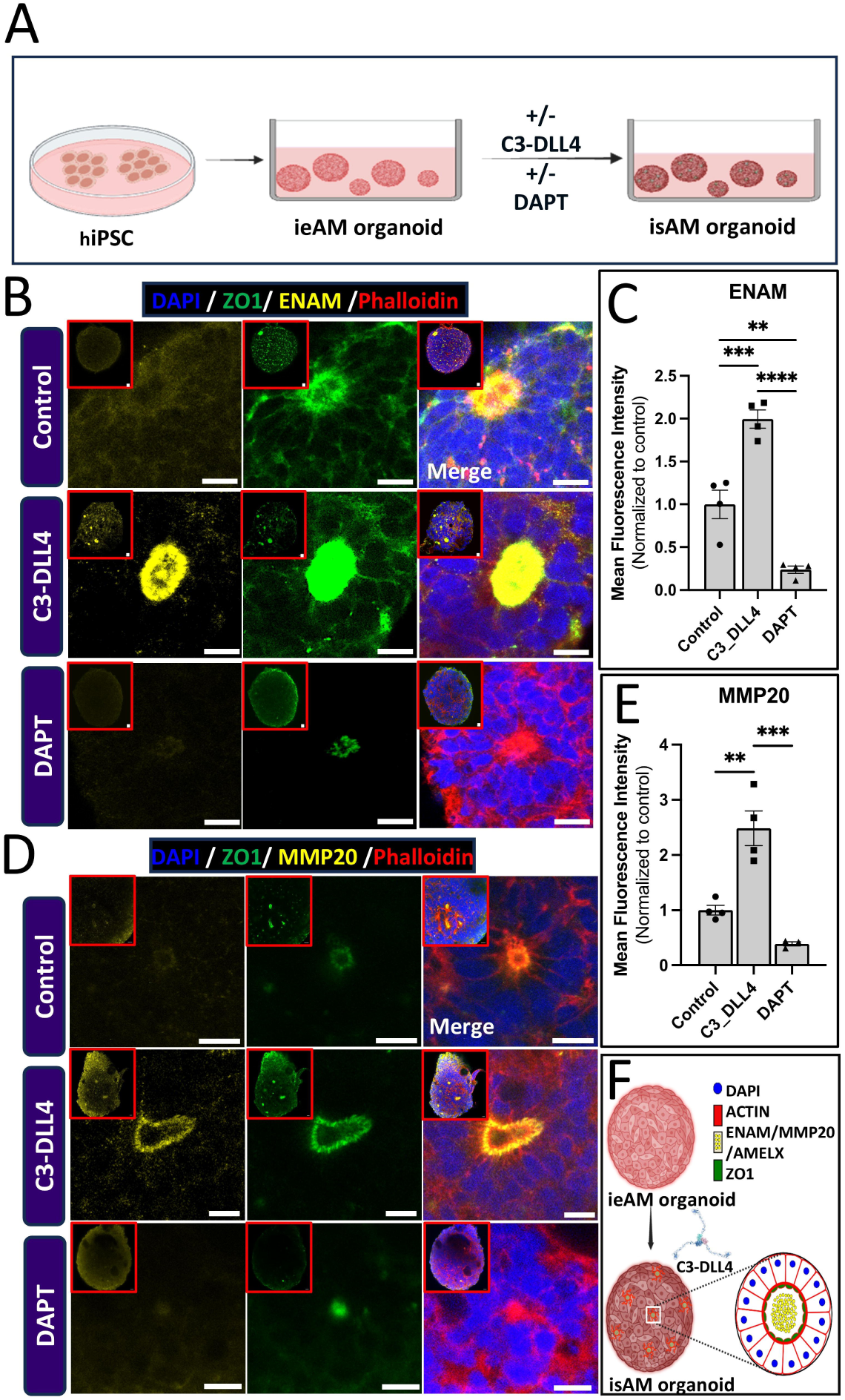
AI-designed Notch activator C3-DLL4 accelerates ameloblast maturation independent of odontoblasts. (A) Schematic representation of the experimental setup showing induced early ameloblast (ieAM) organoid formation from human iPSCs. 3D ieAM organoids are treated with C3-DLL4 (Notch activator) or DAPT (Notch inhibitor), leading to their differentiation into induced secretory ameloblasts (isAM). (B) Immunofluorescence staining of ieAM-derived organoids treated with Control (top), C3-DLL4 (middle), and DAPT (bottom), showing ENAM (yellow) and ZO-1 (green) expression. C3-DLL4-treated organoids display higher ENAM levels and a well-organized structure, whereas DAPT-treated organoids exhibit significantly reduced ENAM expression. Merged images also include DAPI (nuclei, blue) and Phalloidin (cytoskeleton, red). (C) Quantification of ENAM fluorescence intensity, normalized to control, showing a significant increase in C3-DLL4-treated organoids compared to control, while DAPT treatment significantly reduces ENAM expression (\*\*\*\**p < 0.0001*), (***p < 0.001), and (**p < 0.01). (D) Immunofluorescence staining of MMP20 (yellow) and ZO-1 (green) in control, C3-DLL4-, and DAPT-treated organoids. C3-DLL4 enhances MMP20 expression, whereas DAPT treatment drastically reduces it. (E) Quantification of MMP20 fluorescence intensity, demonstrating a significant increase in C3-DLL4-treated organoids (\*\**p < 0.01*) and a significant reduction in DAPT-treated organoids (***p < 0.001). (F) Schematic summary illustrating the transition from ieAM to isAM organoid upon C3-DLL4 treatment, with the upregulation of ENAM and MMP20 and the establishment of an organized, polarized structure.

The treatment of 3D ieAM organoids with C3-DLL4 upregulated expression of key ameloblast maturation markers, including *Enamelin (ENAM)*, *MMP20* and *AMELX* (Figure 3B-3F; Figure S3). This upregulation is indicative of the transition of ieAM organoids to a more mature, secretory ameloblast state (isAM). Notably, the C3-DLL4-treated organoids also displayed enhanced cellular organization and polarization, which are characteristic features of ameloblast maturation in vivo. Since the ameloblast maturation to isAM stage was achieved in the absence of odontoblasts, we conclude that the C3-DLL4 scaffold-based activation of Notch is sufficient to mature ameloblasts to the secretory stage. Furthermore, these data show that soluble C3-Dll4 can substitute for the natural Delta ligand typically provided by iOBs.

Conversely, to evaluated if Notch signaling is essential for iAM maturation in the absence of iOB co-culture, we employed again DAPT, a γ-secretase inhibitor^21^ that prevents Notch receptor activation by blocking the release of the Notch intracellular domain (NICD). Treatment of isolated ieAM organoids with DAPT led to a marked reduction in the expression of key ameloblast maturation markers, such as ENAM, MMP20 and AMELX (Figure 3B-F), indicating that autonomous Notch activation is sufficient for AM progression to a secretory ameloblast (isAM) state. Additionally, organoids exposed to DAPT did not display altered cell organization and polarity, further supporting the idea that active Notch signaling is essential for proper ameloblast maturation to secretory stage (Figure 3B-3E).

Our findings show that the soluble Notch agonist can accelerate the differentiation of iAM organoids into mature, secretory ameloblasts in a defined culture condition, without OB co-culture system (Figure 3F), suggesting that the key function of OB is to activate Notch pathway in AM.

### DLX3 transcription factor is required cell autonomously for ameloblast maturation

Mutations in a key transcription factor, DLX3 has been associated with amelogenesis imperfecta^32,33 34^. While it is known that DLX3 transcription factor directly regulates the expression of DSPP in odontoblast differentiation, it is not clear today if, and at what stage DLX3 is essential for Ameloblast differentiation. Since activation of Notch with designed proteins can generate isAM without OB, we now tested if DLX3 is autonomously required in ameloblast lineage. We generated two independent DLX3 knockout (KO) mutant iPSC lines (KO-10 and KO-13) using CRISPR-Cas9 gene-editing technology. DLX3 protein includes transcriptional activation (TA) domains and the homeodomain. We generated stop codons prior to or on the homeodomain sequence resulting in lack of the key homeodomain helix 3 that facilitates the interaction with target DNA (Figure 4A-B).

**Figure 4.**
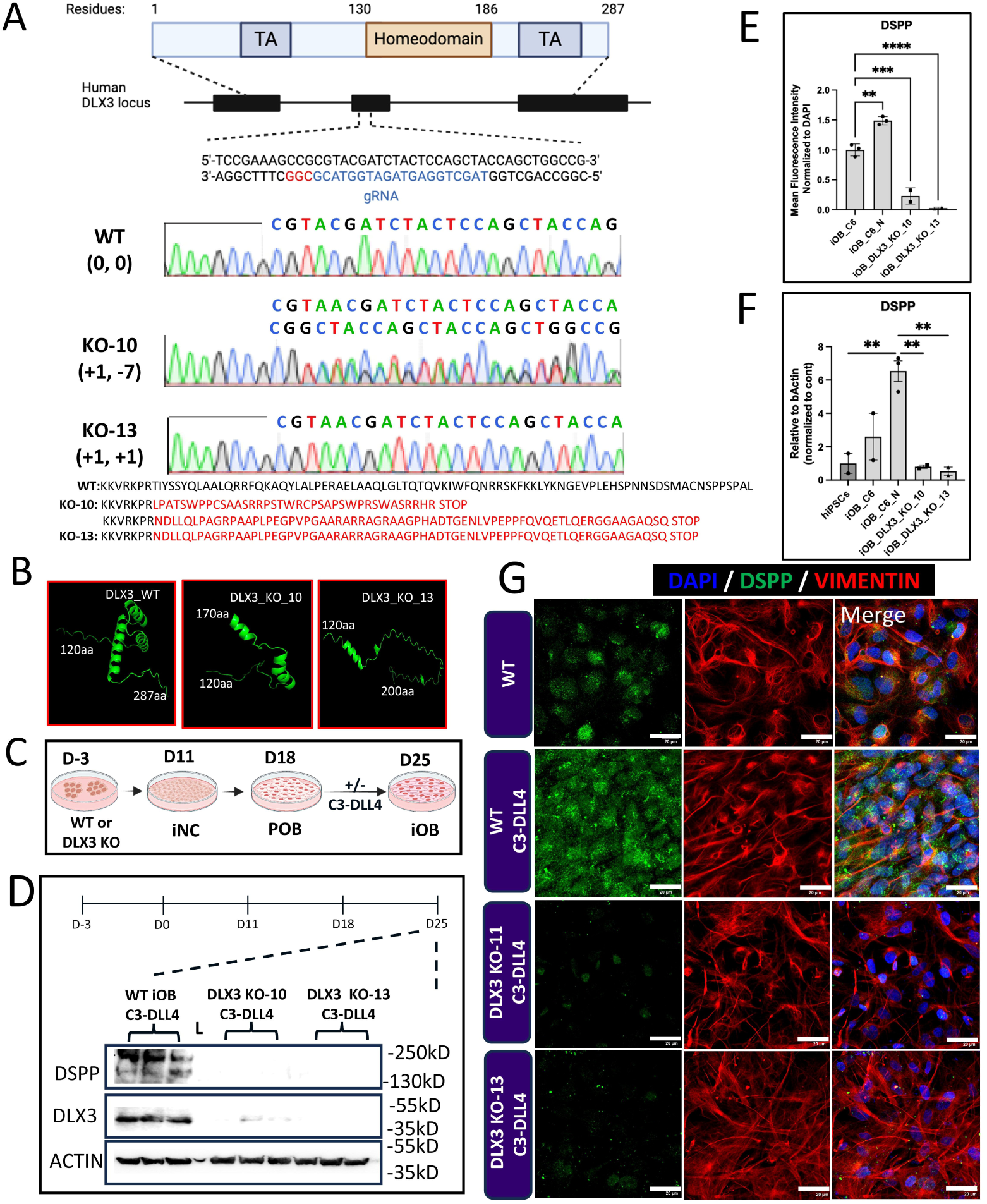
DLX3 is required for odontoblast maturation and DSPP expression. (A) Schematic representation of the DLX3 protein structure and CRISPR-Cas9 targeting strategy. DLX3 consists of two transcriptional activation (TA) domains and a homeodomain essential for DNA binding. The gRNA target site within exon 2 of the human DLX3 locus is shown, along with the corresponding DNA sequence used for gene editing. Below, Sanger sequencing chromatograms compare wild-type (WT) DLX3 with two DLX3 knockout lines (KO-10 and KO-13), revealing distinct insertion and deletion (indel) mutations. KO-10 exhibits a +1 bp insertion and −7 bp deletion, whereas KO-13 carries a +1 bp insertion and −11 bp deletion, leading to frameshift mutations and premature stop codons. The resulting truncated protein sequences confirm loss of the functional DLX3 homeodomain, highlighting its disruption in both knockout clones. (B) Predicted DLX3 homeodomain structures generated using AlphaFold, showing that KO-10 and KO-13 lack critical structural elements compared to WT DLX3. (C) Timeline of odontoblast (iOB) differentiation, showing the differentiation of WT and DLX3 KO iPSCs into neural crest cells (iNCs), pre-odontoblasts (POBs), and mature iOBs. (D) Western blot analysis of DSPP and DLX3 levels in WT and DLX3 KO iOBs. WT iOBs exhibit strong DSPP expression, while DLX3 KO iOBs show a significant reduction in DSPP levels, indicating that DLX3 is required for odontoblast maturation. Actin serves as a loading control. (E) Quantification of DSPP fluorescence intensity from 2D immunofluorescence staining in panel (G), normalized to WT control. WT iOBs treated with C3-DLL4 show a significant increase in DSPP expression, while DLX3 KO iOBs exhibit significantly lower DSPP levels even after C3-DLL4 treatment, reinforcing the requirement of DLX3 for DSPP upregulation (*p < 0.01, ***p < 0.001, \*\*\**p < 0.0001*). (F) qPCR analysis of DSPP expression, normalized to β-Actin, confirming a significant reduction in DSPP transcript levels in DLX3 KO iOBs compared to WT iOBs (p < 0.01), reinforcing that DLX3 is required for odontoblast maturation at the transcriptional level. (G) Immunofluorescence staining of DSPP (green) and Vimentin (red) in WT and DLX3 KO iOBs treated with C3-DLL4. WT iOBs treated with C3-DLL4 maintain DSPP expression, while DLX3 KO iOBs fail to upregulate DSPP, further supporting that DLX3 is essential for odontoblast maturation. Merged images include DAPI (nuclei, blue).

Since DLX3 is known to bind DSPP promoter and control DSPP expression in mature OB^35^, we differentiated DLX3 mutant iOB (Figure 4C). To generate more robust iOB maturation we activated the FGFR1/2c pathway with designed agonist as done before^18^. The previous analysis of human fetal single cell sequencing data predicted Notch pathway function in POB to OB differentiation/maturation^18^. We therefore tested and showed that iOB maturation measured by DSPP expression was markedly increased with Notch activation using designed Notch agonist C3-DLL4 (Figure 4E-G). We differentiated wild type and DLX3 mutant iPSC to mature iOB using the designed agonists. With this matured iOB differentiation assay we observed a dramatic reduction of DSPP in DLX3 mutant OB (compared to wild type), both on protein and RNA level (Figure 4D-G). These data confirm that our DLX3 mutants behave as expected showing significantly reduced expression of DSPP in OB lineage. This further confirms that DLX3 can activate DSPP transcription cell autonomously in mature OB.

To dissect DLX3 function in Ameloblasts, we differentiated the DLX3 knockout (KO) iPSC lines into first induced early ameloblasts in 2D model and then induced secretory ameloblasts (isAM) in 3D organoid model using Notch activator, C3-DLL4 (Figure 5A). Quantitative comparisons using Western blot analysis between differentiated DLX3 KO and wild type-iAM D16 cells revealed that the absence of DLX3 did not affect the expression of early ameloblast markers, including *AMBN, SP6*, or transient DSPP expression (Figure 5B). These findings suggest that DLX3 is not required for the early ameloblast differentiation or regulation of DSPP transcription in ameloblasts. Similarly, when tested in matured isAM organoid system the immunostaining analysis of ZO-1 revealed a normal cellular polarity of DLX3 mutant ameloblasts, with apical lumen marked with ZO-1. However, a significant decrease in the expression of essential maturation markers, such as *ENAM* and *MMP20* was observed in the DLX3 KO organoids compared to controls (Figure 5C-5F). This reduction reveals an essential role of DLX3 in ieAM transition to the mature, secretory ameloblast state.

**Figure 5.**
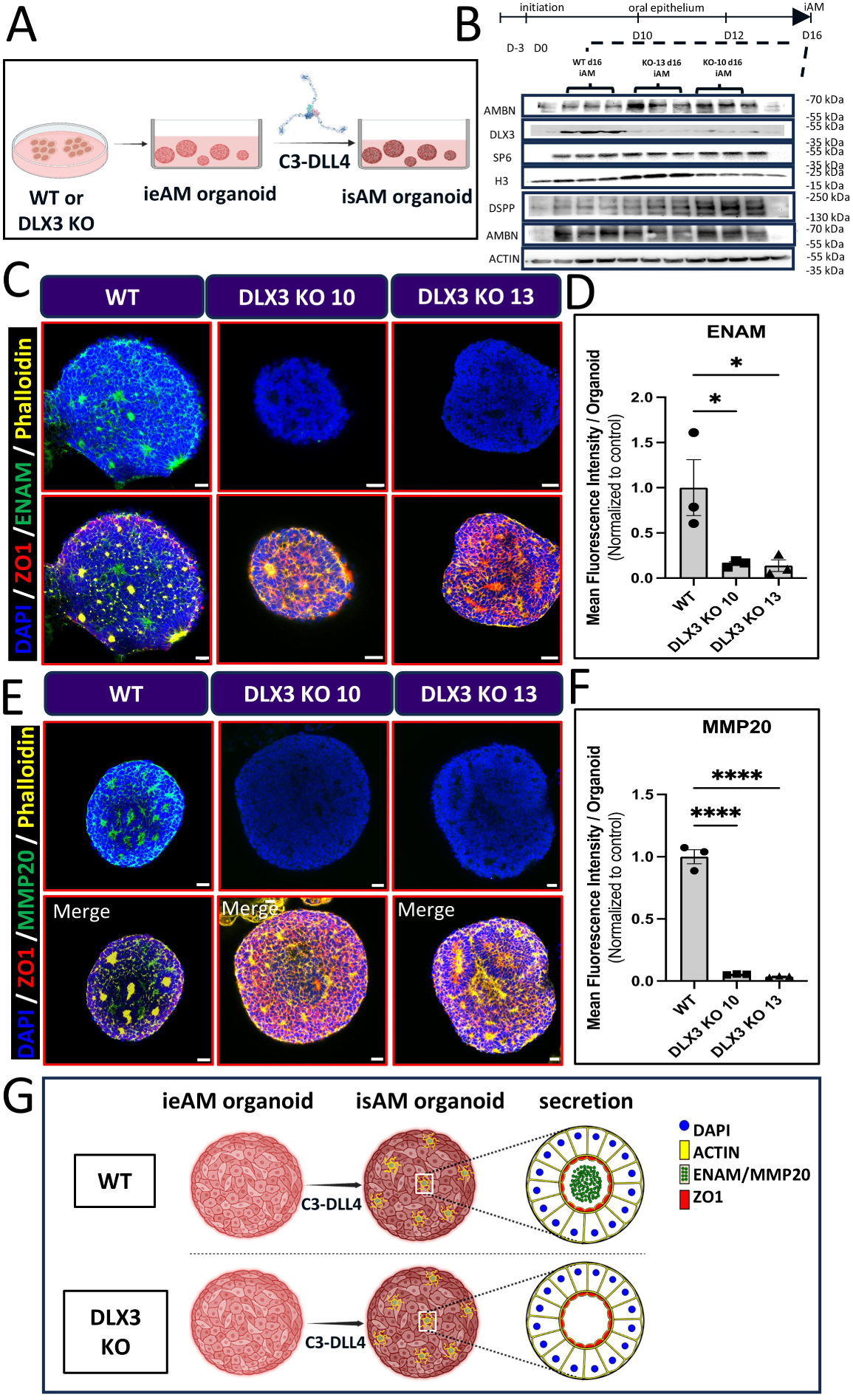
DLX3 is required for ameloblast terminal maturation and enamel matrix protein expression. (A) Schematic representation of the experimental setup, where wild-type (WT) or DLX3 knockout (KO) iPSCs are differentiated into induced early ameloblast (ieAM) organoids and further matured into induced secretory ameloblasts (isAM) organoids upon C3-DLL4 treatment. (B) Western blot analysis of AMBN, DLX3, SP6, H3, and DSPP protein levels in WT and DLX3 KO iAM organoids at day 16 (d16). DLX3 KO iAM organoids show significantly reduced expression of enamel matrix proteins, indicating a defect in ameloblast maturation. ACTIN serves as a loading control. (C) Multi-stack confocal images and Immunofluorescence staining of ENAM (green) and ZO-1 (red) in WT and DLX3 KO organoids. WT organoids show strong ENAM expression, whereas DLX3 KO organoids exhibit markedly reduced ENAM levels. Merged images also include DAPI (blue, nuclei) and Phalloidin (yellow, cytoskeleton). (D) Quantification of ENAM fluorescence intensity, showing a significant reduction in DLX3 KO organoids compared to WT (\**p < 0.05*). (E) Multi-stack confocal images and Immunofluorescence staining of MMP20 (green) and ZO-1 (red) in WT and DLX3 KO organoids. WT organoids exhibit high MMP20 expression, while DLX3 KO organoids show dramatically reduced MMP20 levels. (F) Quantification of MMP20 fluorescence intensity, confirming a significant decrease in DLX3 KO organoids (\*\*\*\**p < 0.0001*). (G) Schematic model summarizing the impact of DLX3 KO on ameloblast maturation, showing that WT isAM organoids display correct polarization and secretion of enamel matrix proteins (ENAM/MMP20), whereas DLX3 KO organoids fail to properly mature.

Our findings demonstrate that while DLX3 is not necessary for early ameloblast differentiation or polarity establishment, it is crucial for the terminal maturation phase, particularly in driving the expression of enamel matrix proteins (Figure 5G). These data show that DLX3 is required on cell autonomous level for late ameloblast maturation.

## Discussion

We identified the Notch pathway in the interaction between odontoblasts (OB) and ameloblasts (AM), which leads to AM maturation. The expression patterns of Notch and Delta genes supported the hypothesis that Delta from OB can activate Notch in AM. To test whether Notch pathway activation is necessary and sufficient for AM maturation, we utilized small molecules and novel AI-designed protein scaffolds. Our results indicate that Notch inhibition (using DAPT) affects enamelin (ENAM) secretion in co-culture, demonstrating that Notch is required for AM maturation. Next, we utilized our designed soluble Notch scaffold that can activate Notch processing presumably due to the forces generated by the scaffold interacting with Notch in the surface of neighboring cells. Notch activation with this soluble agonist enhanced the expression of mature isAM markers, ENAM, MMP20 and AMELX, even in the absence of co-culture with OB cells. These data suggest that Notch pathway activation is critical for AM maturation, and the designed Delta-scaffold can substitute for OB in this process. This newly developed maturation paradigm allowed us to reveal that the Amelogenesis Imperfecta gene DLX3 is required not only in OB but also in the AM cell lineage. Specifically, DLX3 is not necessary for the pre-ameloblast stage, but it is essential for the expression of ENAM and MMP20 during the ameloblast maturation stage.

In this paper we utilized a designed scaffold that leverages the natural ligand-receptor interaction mechanism of Notch pathway by incorporating engineered extracellular domain of DLL4 tethered to computer designed scaffold protein. This design ensures stable ligand presentation and facilitates efficient engagement with Notch receptors on target cells. Moreover, the scaffold allows for precise spatial positioning and temporal control over Notch activation. By mimicking native ligand dynamics in a controlled environment, this system provides unparalleled flexibility to dissect the nuances of Notch signaling. Importantly, it enabled investigation of how spatially and temporally regulated Notch signaling orchestrates the complex processes underlying ameloblast differentiation and maturation.

Potential crosstalk between ameloblasts and odontoblasts has been proposed previously. Although an intact basement membrane typically separates the two cell types, this barrier disintegrates at the onset of predentin secretion, providing a narrow time frame for direct cell– cell interactions^31^. We have now revealed a plausible mechanism for this interaction that involve Notch ligands (e.g., Jagged or Delta), which may become accessible to ameloblast processes upon basement membrane breakdown. Further investigation of this Notch signaling pathway, particularly through advanced imaging techniques, could yield critical insights into how odontoblasts communicate with ameloblasts to coordinate dentin and enamel formation.

DLX3 has been shown to positively regulate the expression of crucial enamel matrix proteins, including AMELX, ENAM, and ODAM. Although AMBN has not been as extensively studied, it is believed that DLX3 also influences AMBN expression as part of its broader role in the enamel gene network. Mutations or reduced expression of DLX3 have been associated with enamel defects, suggesting potential alterations in AMBN expression. However, it remains unclear which of these phenotypes are cell-autonomous and which result from the interactions between odontoblasts (OB) and ameloblasts (AM). In this study, we reveal that DLX3 is specifically required for the terminal maturation phase of ameloblasts in a cell-autonomous manner, particularly in driving the expression of enamel matrix proteins, independent of odontoblast contribution. Our work to generate matured isAM organoids without the co-culture of odontoblasts has enabled a detailed dissection of DLX3’s role in ameloblast differentiation. This study clarifies that DLX3 functions cell autonomously in AM to support proper enamel generation and raises questions about DLX3’s direct targets in AM.

Our innovative protein design approach to activate Notch has yielded critical insights into the molecular mechanisms of Notch signaling during enamel organ development and facilitated the generation of a more advanced platform. This understanding has deepened our knowledge of Amelogenesis Imperfecta and will pave the way for future research aimed at developing therapies.

### Limitations of the Study

Our study primarily relies on iPSC-derived models from a select group of cell lines, which has allowed us to achieve robust reproducibility; however, expanding the analysis to include additional iPSC sources could further validate the generalizability of our findings. Additionally, while our data suggest that Notch activation is sufficient to bypass odontoblast co-culture, additional studies may provide deeper insights into how closely the in vitro signaling dynamics reflect the complex in vivo microenvironment during enamel formation.

## Resource availability

### Lead contact

Further information and requests for resources and reagents should be directed to and will be fulfilled by the lead contact, Hannele Ruohola-Baker.

### Materials availability

This study did not generate new unique reagents.

### Data and code availability

The lead contact will share all data analyzed and reported in this paper upon request. This paper does not report the original code. Any additional information required to reanalyze the data reported in this work is available from the lead contact upon request.

## Acknowledgments

We thank the Ruohola-Baker lab members for their helpful discussions. We thank Khushal Thakor, Emma D Cox, Leah Tadese, Chris Cavanaugh, Jennifer Hesson, and Yen C Lim for their technical assistance, Dr. Yan Ting Zhao for help with molecular biology, and Dale Hailey and the Garvey microscopy core for help with microscopy. This work is supported by ISCRM Fellows Program (Anjali Patni) and grants from the National Institutes of Health DE033016 (J.M., H.R.-B), 1P01GM081619, R01GM097372, R01GM083867, NHLBI Progenitor Cell Biology Consortium (U01HL099997; UO1HL099993) SCGE COF220919 (H.R-B), and AHA 19IPLOI34760143, Brotman Baty Institute (BBI), DOD PR203328 W81XWH-21-1-0006 and Stem Cell Gift Funds for H.R-B. The Birth Defects Research Laboratory was supported by NIH award number 5R24HD000836, to IAG and DD, from the Eunice Kennedy Shriver National Institute of Child Health and Human Development.

## Author contributions

Conceptualization: A.P.P., J.M., and H.R.-B.; A.P.P. and H.R.-B.; conceived and analyzed the designed proteins in ameloblast and odontoblast differentiation; Methodology: A.P.P., A.A., R. Mout, R. Moore, J.M., and H.R.-B.; investigation: A.P.P., A.A.; A.P.P. performed organoid assays; funding acquisition: A.P.P., H.R.-B., and J.M.; resources: R. Mout, G.Q.D., and H.R.-B.; supervision: R. Mout, H.R.-B., and J.M.; visualization: A.P.P., A.A. and H.R.-B.; A.P.P., R. Mout, A.A., and R. Moore prepared the figures; formal analysis: A.P.P., A.A.; data curation: A.P.P.; project administration: H.R.-B.; writing – original draft, A.P.P., A.A., H.R.-B, and J.M.; writing – review and editing, A.P.P., R. Mout, R. Moore, A.A., H.R.-B. and J.M., G.Q.D., D.B.

## Supplemental Figure Legends

**Figure S1.**
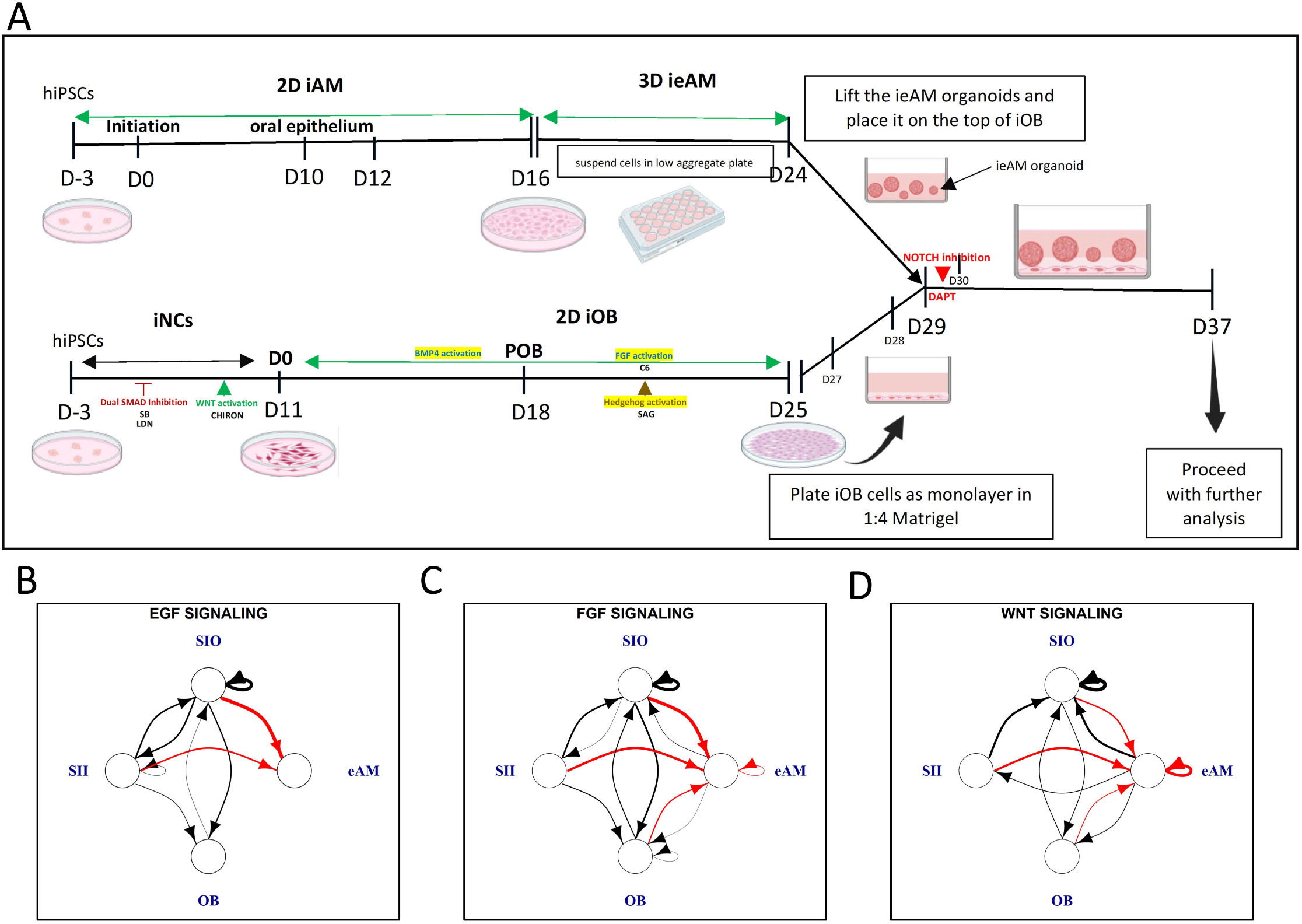
Schematic overview of the co-culture differentiation timeline and signaling interactions in ameloblast maturation. (A) Timeline of the ameloblast-odontoblast co-culture system, illustrating the differentiation of human iPSCs into induced early ameloblasts (ieAM) and induced odontoblasts (iOB). hiPSCs were first differentiated into iNCs using dual SMAD inhibition (SB, LDN) and WNT activation (CHIR). iNCs were further differentiated into iOBs using BMP4, FGF (C6), and Hedgehog activation (SAG). Simultaneously, iAM differentiation followed a 2D culture phase until Day 16, after which cells were transferred to ultra-low attachment plates to generate 3D ieAM organoids. On Day 24, ieAM organoids were placed onto the iOB monolayer to establish the co-culture system. Notch inhibition (DAPT) was introduced at Day 29 for 24 hours, followed by a media change on Day 30 to evaluate its effect on ameloblast maturation. The co-culture was maintained until Day 37, when samples were collected for analysis. (B-D) Predicted signaling interactions in the co-culture system based on pathway analysis. (B) EGF signaling, (C) FGF signaling, and (D) WNT signaling illustrate intercellular communication between OBs and eAMs, with red arrows representing predominant incoming signals to eAMs. Black arrow indicates other possible interaction not considered in the analysis of TopPath. The inner stratum intermedium (SII) and outer stratum intermedium (SIO) cell populations are included, suggesting additional signaling interactions in ameloblast maturation.

**Figure S2.**
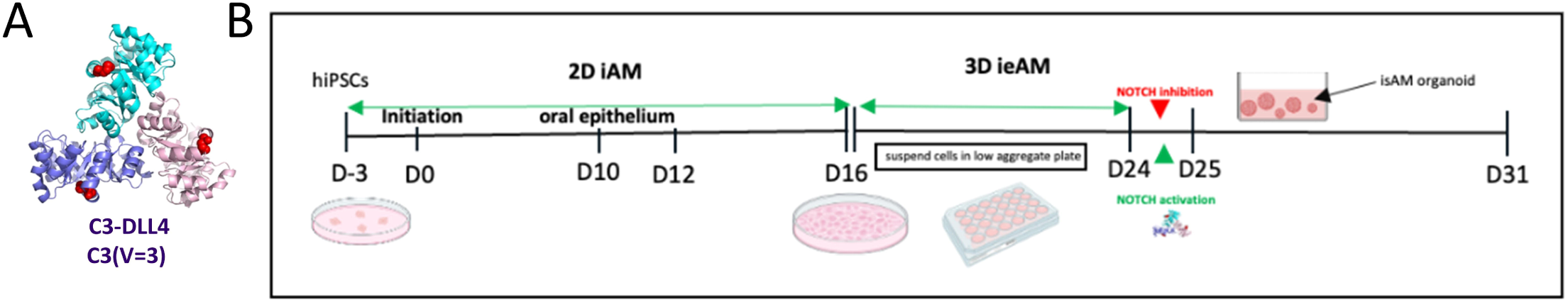
Notch activation and inhibition in ameloblast differentiation. (A) Structural representation of C3-DLL4, a computationally designed trimeric scaffold conjugated to DLL4, used to activate Notch signaling during ameloblast differentiation. (B) Timeline of iPSC-derived ameloblast differentiation, illustrating the transition from 2D iAM culture to 3D ieAM organoid formation. hiPSCs were differentiated into 2D iAMs from Day 0 to Day 16, after which cells were suspended in low-attachment plates to generate 3D ieAM organoids. Notch activation was induced on Day 24 using C3-DLL4 for 24 hours. Similarly, Notch inhibition was introduced on Day 24 for 24 hours, followed by a media change on Day 25. The organoids were kept until Day 31 to evaluate its effect on ameloblast maturation.

**Figure S3.**
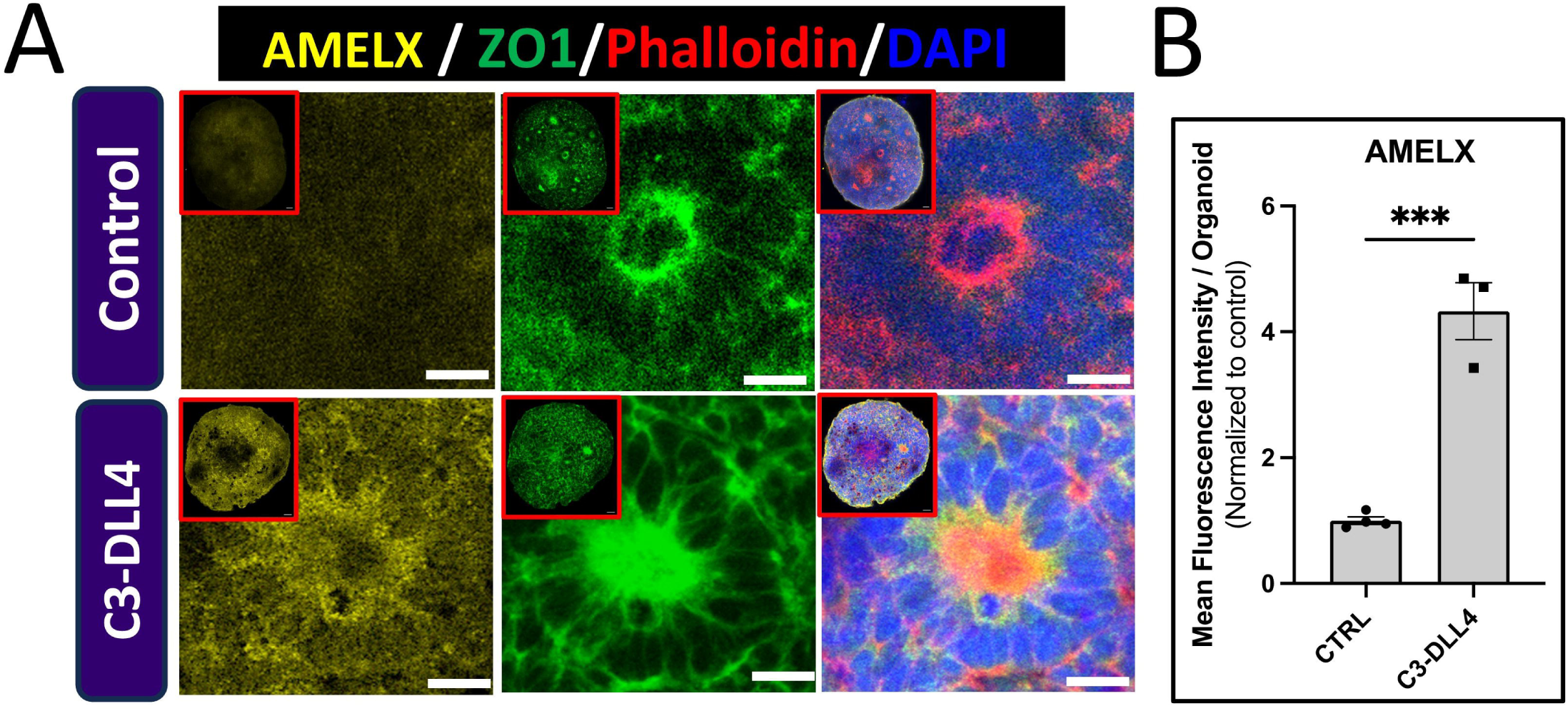
Notch activation by C3-DLL4 enhances AMELX expression in ameloblast organoids. (A) Multi-stack confocal images and Immunofluorescence staining of AMELX (yellow), ZO-1 (green), Phalloidin (red, cytoskeleton), and DAPI (blue, nuclei) in induced early ameloblast (ieAM) organoids treated with either control conditions (top) or C3-DLL4 (bottom). C3-DLL4-treated organoids exhibit significantly increased AMELX expression compared to controls, with enhanced epithelial organization and polarity. Insets highlight representative individual cells within the organoids. (B) Quantification of AMELX fluorescence intensity per organoid, normalized to control conditions. C3-DLL4 treatment significantly increases AMELX expression compared to controls (***p < 0.001).

**Figure S4.**
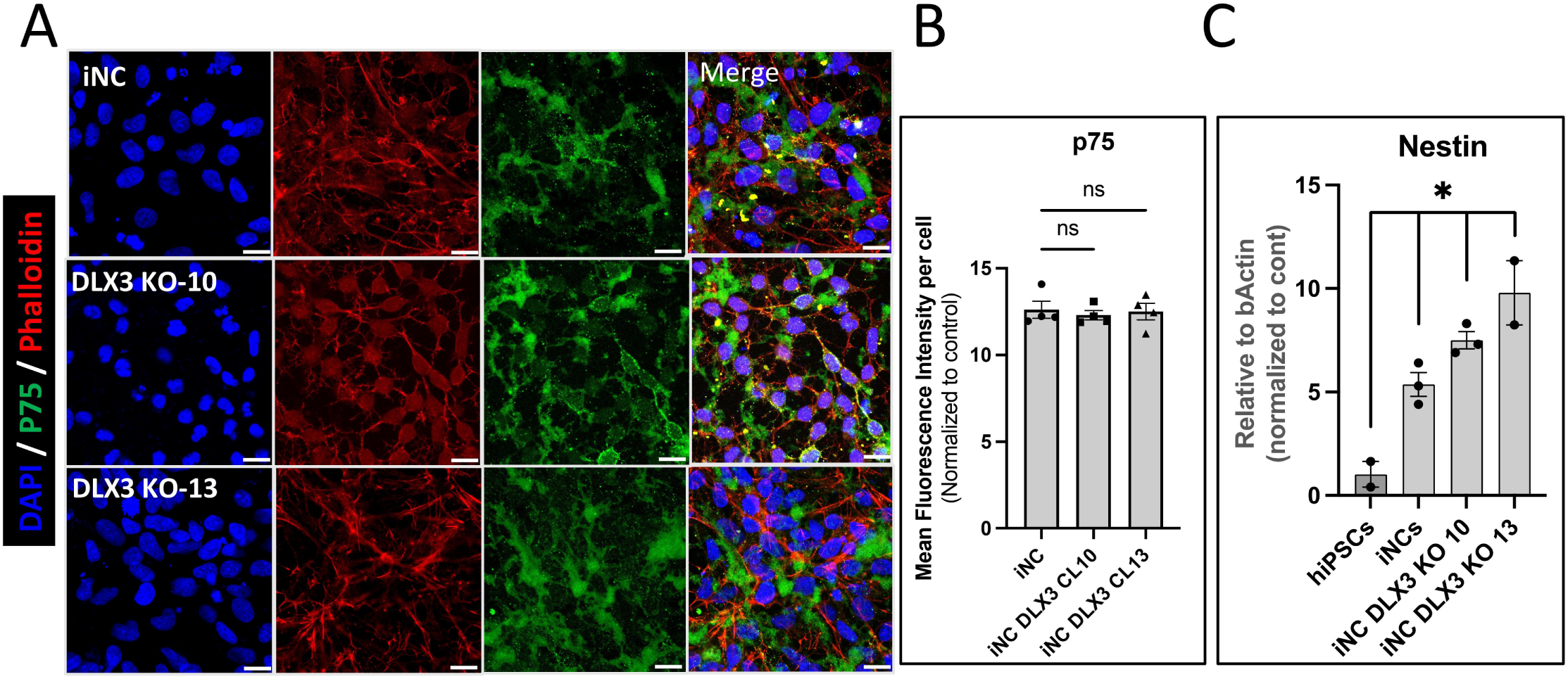
DLX3 knockout does not alter neural crest (iNC) identity. (A) Immunofluorescence staining of p75 (green), Phalloidin (red, cytoskeleton), and DAPI (blue, nuclei) in WT and DLX3 knockout (KO-10, KO-13) iNCs. All conditions show comparable p75 expression, indicating that DLX3 knockout does not impair iNC identity. Merged images highlight cellular morphology and cytoskeletal organization. (B) Quantification of p75 fluorescence intensity per cell, normalized to control iNCs. No significant differences (ns) are observed between WT and DLX3 KO iNCs, confirming that p75 expression remains unaffected by DLX3 loss. (C) qPCR analysis of Nestin expression, normalized to hiPSC, showing no significant difference between WT and DLX3 KO iNCs, indicating that Nestin transcription expression remains unchanged upon DLX3 loss.

## Materials and Methods

### iPSC derived Ameloblast differentiation

Briefly, hiPSCs (WTC-11 human induced pluripotent stem cells) (Coriell, #GM25256) were seeded on 12-well plates coated with growth factor-reduced Matrigel (Corning, #356231) and cultured in mTeSR medium (StemCell Technologies, #85850) until cells reach confluency with medium changes daily. On the first day of differentiation (Day 0), stem cell media is replaced with base iAM media consisting of EpiCult-C media (StemCell Technologies, #05630) supplemented with 1X EpiCult-C Proliferation Supplement (STEMCELL Technologies), 0.1uM β-mercaptoethanol (BME) (Sigma, #M7522), 1X Hydrocortisone Stock Solution (96 μg/mL), 0.05X GlutaMAX (Gibco), 0.4X NEAA (Gibco) and 1X Penicillin-Streptomycin (Gibco) and 400nM smoothened agonist (SAG) (Selleckchem, # S7779). On day 3 of differentiation 150pM of monomeric bone morphogenic protein-4 (BMP4) (rndsystems, #314-BP-010) is added daily until day 7. At day 8, the base media is supplemented with 1uM of BMP-I inhibitor (LDN-193189) (Tocris, #6053), 5uM of GSK3-Inhibitor (CHIR99021) (Selleckchem, #4423), 500pM epidermal growth factor (EGF) (rndsystems, #236-EG) and 70pM of Neurotrophin-4 (NT4) (rndsystems, #268-N4). No media refresh is added on day 9. On day 10, repeat with the same small molecules used on day 8. From day 12 onwards the cultures were then extended until day 16 by adding 1X N2 Supplement (Gibco #17502-048), 400nM SAG, 300pM BMP4, 5uM of GSK3-Inhibitor (CHIR99021), 500 pM of EGF 2nM transforming growth factor beta 1(TGFβ1) (rndsystems, #7754-BH) 70pM of Neurotrophin-4 (NT4) (rndsystems, #268-N4) and 300pM of Activin (Peprotech, #120-14P) and for the early ameloblast stage at day 16. The media containing the small molecules were changed every other day from day 12 until day 16.

### Production and Characterization of the C3-DLL4 Notch Activator

The C3-DLL4 Notch activator was produced using a previously established computationally designed scaffold with defined valency and geometry. The C3 scaffold, a homotrimeric helical bundle, was conjugated to DLL4 using SpyLigation, which forms a stable isopeptide bond between SpyCatcher (SC) and SpyTag (ST). The C3-SpyCatcher fusion was expressed in *E. coli*, while DLL4-SpyTag was produced in mammalian Expi293F cells, ensuring proper folding and post-translational modifications. Both components were purified using affinity chromatography, and successful conjugation was confirmed by SDS-PAGE and Coomassie staining, showing a higher molecular weight band corresponding to the assembled C3-DLL4 complex. For cell treatment, a 10 µM stock solution of C3-DLL4 was prepared by conjugating DLL4-ST with C3-SpyCatcher, followed by serial dilutions to achieve precise ligand presentation and receptor engagement.

### Notch Activation assay in reporter cell line

The Notch activation assay was performed using U2OS-N1-Gal4/UAS-H2B-mCitrine cells^36,37^ to evaluate the effect of C3-DLL4 on Notch signaling. Cells were seeded in a 96-well tissue culture-treated plate at a density of 15,000 cells per well in DMEM supplemented with 10% FBS, 1X Penicillin-Streptomycin, 1X GlutaMax, and 1X Sodium Pyruvate. Doxycycline (Dox, 1 µg/mL) was added at seeding to induce Notch1-Gal4 (N1-Gal4) expression, which is essential for the assay. On Day 1, cells were treated with C3-DLL4. Through serial dilutions of C3-DLL4 the final concentrations of 200 nM, 100 nM, 50 nM, 10 nM and 1nM (DLL4 molar equivalent) were added to respective wells in replicates. Untreated wells served as negative controls. After treatment, the cells were gently mixed and incubated under standard culture conditions at 37°C with 5% CO□. On Day 2, the media was replaced with fresh Dox-containing media to maintain Notch induction. On Day 4, the Notch activation was assessed by measuring H2B-mCitrine fluorescence. Fluorescence intensity was quantified to determine the dose-dependent activation of Notch signaling. Control wells without C3-DLL4 were analyzed to establish baseline fluorescence, ensuring accurate interpretation of Notch activation levels.

### *iPSC* derived Odontoblast differentiation

We modified the odontoblast differentiation protocol previously described by our lab^3,18^ to avoid serum usage. This differentiation protocol begins with hiPSCs, treated with dual SMAD inhibitors, SB431542 and LDN-193189 (Tocris, #6053), to inhibit TGF-β and BMP signaling pathways, respectively. This promotes ectodermal lineage commitment, guiding the hiPSCs toward an odontogenic trajectory. From Day 0 to Day 11, the WNT signaling pathway is activated using CHIR99021, a GSK-3β inhibitor, to direct mesenchymal lineage specification and to facilitate the generation of neural crest cells (iNCs), a precursor population essential for odontoblast development. On Day 12, BMP4 is introduced to drive the differentiation of iNCs into odontoblast precursors in serum-free conditions.

Specifically, p75□+□iNC cells were cultured in Serum-free Odontogenic Medium, consisting of DMEM□+□Glutamax (Gibco 10566016), 100 nM dexamethasone (Sigma-Aldrich D4902), 15% KnockOut Serum Replacement (KOSR), 0.005mM ITS-A (Gibco 51300-044), 5 mM β-glycerophosphate (Sigma-Aldrich G9422), and 50 µg/ml L-ascorbic acid (Sigma-Aldrich A4544) for 14 days (OB). Odontogenic Medium was supplemented with 50 ng/ml BMP4 (Stemcell Technologies 78211) for Day 11 to Day 18, followed by 25 ng/ml BMP4 (Stemcell Technologies 78211) and 400 nM SAG (Stemcell Technologies 73412) from Day 18 until Day 25 supplemented with 100 ng/ml C6^18,20^ for 14 days (iOB C6); followed by treatment with or without Notch activator (50 nM C3-DLL4)^17,18^ (iOB C6 N) on Day 18 for 24 hours followed by change of media on Day 19 followed until Day 25. All cultures were performed on Matrigel-coated plates at a 1:30 dilution and incubated at 37°C with 5% CO2. Each differentiation was performed in triplicate, with undifferentiated hiPSC as the negative control.

### Establishing a Functional Co-Culture model for ameloblast and odontoblast

Human induced pluripotent stem cells (hiPSCs) were differentiated into ameloblast and odontoblast lineages to establish a functional co-culture model for ameloblasts and odontoblast lineages under defined serum-free conditions. For ameloblast differentiation, hiPSCs were cultured in a 2D monolayer to generate early ameloblasts (ieAM) from Day 0 to Day 16. On Day 16, differentiated iAM cells were trypsinized using TrypLE Express (Thermo Scientific or Gibco, #12604013). The cells were gently resuspended and then transferred into 24-well ultra-low attachment plates for three-dimensional (3D) suspension culture in Epicult Plus medium (Stemcell Technologies 06070), allowing them to self-organize into ieAM organoids. The ieAM organoid cultures were incubated at 37^°^C with 5% CO_2_, and the media was changed every other day. These organoids were cultured until the next 7 days, stabilizing their maturation. Concurrently, odontoblast cells were made similarly in a separate plate by culturing iOBs differentiated from iNCs (neural crest) that were differentiated from hiPSCs in a serum-free odontogenic differentiation medium. The differentiated iOB cells were plated as monolayer mixed in 25% (v/v) of Matrigel (Corning, #356231) diluted in odontogenic media in a glass-bottomed 24-well plate (Corning, #3603). The next day, ieAM organoids suspended in the ameloblast base medium and 10 μM ROCKi (Y-27632, Selleckchem, #S1049) were added on top of the iOB monolayer and then incubated for 24 hours at 37°C in 5% CO_2_. The co-culture was supplemented with fresh media (1:1 mixture of ameloblast and odontogenic media) every three consecutive days. The co-culture was collected on Day 37 for further analysis.

### DAPT inhibition assay

To assess the effect of Notch inhibition on ameloblast-odontoblast interactions, a DAPT inhibition assay was performed after 4 days of the co-culture (Day 29). DAPT (10 μM) (SantaCruz Biotech #sc-201315) was added to the co-culture medium to inhibit Notch signaling, and the treated co-culture was incubated for 24 hours at 37^°^C with 5% CO□. After 24 hours, the media was replaced with a fresh co-culture medium to continue differentiation. The co-culture was maintained under standard conditions and collected on Day 37 for further analysis of ameloblast maturation and enamelin secretion.

### Notch Activation in the Co-Culture Assay Using C3-DLL4

Notch activation was induced in the co-culture system by treating ieAM organoids with C3-DLL4 (50 nM) after 4 days of co-culture on Day 29. The treatment was maintained for 24 hours, after which the media was replaced on Day 30. Co-cultures were maintained with media changes every three days until Day 37, when they were collected for further analysis.

### Pathway analysis (TopPath)

To reanalyze the pathway involved in ameloblast maturation, we used TopPath pipeline, as described previously with minor changes. In brief, the talklr R package^19^ was used to pinpoint ligand-receptor interactions specific to each cell type during the transition stage of ameloblast maturation from early ameloblast to secretory ameloblast. The DEsingle^38^ and scMLnet^39^ tools were applied to analyze downstream signaling by creating multilayer networks that connect ligands to receptors and transcription factors to their corresponding differentially expressed target genes. Pathway activity scores were then calculated, reflecting the percentage (0-100%) of total activity across all pathways evaluated in the analysis. The pathways considered in the analysis were: TGFβ, BMP, GDF, GDNF, NODAL, ACTIVIN, WNT, ncWNT, EGF, NRG, FGF, PDGF, VEGF, IGF, INSULIN, HH, EDA, NGF, NT, FLT3, HGF, NRXN, OCLN, NOTCH.

### Generation of DLX3 KO iPSC

One million WTC11 iPSC were electroporated with Cas9 (0.3uM, Sigma) and gRNA targeting DLX3 (1.5uM, Synthego) as RNP complex using Amaxa nucleofector (Human Stem Cell kit 2) in presence of ROCK inhibitor. Individual colonies were hand-picked and plated into 96 well plates. DNA was extracted using Quick Extract DNA extraction solution (Epicentre #QE09050) and nested PCR was performed using Phusion Flash polymerase (ThermoFisher, #F631S). The PCR product was purified using ExoSap-IT (Thermofisher) and sent for Sanger sequencing analysis (Genewiz, Azenta Life Sciences) to identify potential KO clones. gRNA sequence: TAGCTGGAGTAGATCGTACG. PCR primer sequences: F: GAAGGCGTCGTGAGCGAAG, R: TAGCCTGGAGGGAAAACACG. KO was verified at the protein level after 14 days of odontoblast differentiation and 16 days of ameloblast differentiation.

### Notch Activation and Inhibition in Ameloblast organoid Differentiation

hiPSCs were differentiated into 2D induced ameloblasts (iAMs) from D0 to Day 16. At Day 16, iAMs were transferred to low-attachment plates to form 3D induced early ameloblast (ieAM) organoids in Epicult Plus medium (Stemcell Technologies #06070). Notch activation was induced at Day 24 using the C3-DLL4 Notch activator (50nM), a trimeric scaffold conjugated to DLL4, or it was incubated with Notch inhibitor, DAPT (10 μM) (SantaCruz Biotech #sc-201315) (Figure A), for 24 hours. On Day 25, the media was replaced, and the ieAM organoids continued maturing into induced secretory ameloblasts (isAMs) by Day 31 (Figure B). Likewise, DLX3 knockout clones (Clone 10 and Clone 13) were differentiated following the same protocol.

### Immunostaining and Confocal Imaging

For immunostaining, the organoids were fixed in 4% paraformaldehyde (PFA), then immersed in 1X PBS for 3x 5-minute washes, and then immersed in 0.5% TritonX 100 at RT for 10 minutes to facilitate permeabilization. Later, Organoids were blocked in solution (1X PBS containing 10% BSA, 5% normal goat serum, and 0.1% Triton-X) for 2 hours. All incubations were done in tubes on a nutator. They were then suspended in antibody dilution buffer (1X PBS containing 10% BSA, 5% normal goat serum, and 0.2% Triton-X) and primary antibodies diluted at concentrations recommended by the manufacturer and incubated overnight at 4°C. On the second day, cells were washed 3x for 6 minutes with 1X PBS, resuspended in antibody dilution buffer containing secondary antibodies at 1:200 and DAPI (1:50), and then incubated overnight at 4°C. On the third day, organoids were washed 3x for 6 minutes in 1x PBS and mounted in Vectashield Antifade Mounting Media on a glass concavity microscope slide, one to three organoids per well. Organoids were then imaged using a Leica (DMi8) SP8 LIGHTNING confocal microscope (25x and 40x objectives) equipped with HyD and PMT spectral detectors and Leica LASX acquisition software [version 3.5.5IR], or a High Resolution Widefield Nikon ECLIPSE Ti Fluorescent microscope and Nikon Ti2 confocal microscope with Nikon NIS-Elements software.

### Quantification of Immunofluorescence Signals

Immunofluorescence signals for the proteins of interest (ENAM, AMELX, and MMP20) were quantified as follows. First, raw images were processed using Fiji (ImageJ v2.3.091/92) to split each image stack into individual TIFF files corresponding to DAPI and the target protein. To normalize quantification across each organoid, nuclei were segmented using the StarDist3D pipeline on ZEISS arivis Cloud (Apeer) and then counted with ImageJ’s 3D Object Counter, ensuring that threshold parameters remained consistent across images acquired under similar conditions.

The fluorescent signals for the proteins of interest were quantified in three dimensions by segmenting puncta with the ImageJ 3D Object Counter. The ImageJ measure function was then used to determine both the mean fluorescence intensity and the integrated intensity for each punctum. The overall mean signal for each organoid was normalized to the nuclei count. Finally, all results were compiled into a spreadsheet and analyzed with GraphPad Prism to assess the statistical significance between groups and conditions.

### RNA Extraction and QRT-PCR Analysis

RNA was isolated from the cells utilizing Trizol reagent (Life Technologies) in accordance with the manufacturer’s guidelines. RNA samples underwent treatment with Turbo DNase (Thermo Fisher Scientific) to eliminate genomic DNA contamination and were subsequently quantified utilizing the Nanodrop ND-1000 spectrophotometer (Thermo Fisher Scientific). For cDNA synthesis, 1□µg of RNA was reverse transcribed using the iScript™ cDNA Synthesis Kit (Bio-Rad) or the Applied Biosystems™ High-Capacity cDNA Reverse Transcription Kit (Thermo Fisher Scientific). Quantitative real-time PCR (qPCR) was performed on 10□ng of cDNA per reaction using SYBR Green (Applied Biosystems) and the 7300 Real-Time PCR System (Applied Biosystems). The conditions for the polymerase chain reaction (PCR) were established as follows: an initial incubation at 50□°C for 2 minutes, followed by a denaturation step at 95□°C for 10 minutes. This was succeeded by 40 amplification cycles, each consisting of a denaturation phase at 95□°C for 15 seconds and an annealing/extension phase at 60□°C for 1 minute. All quantitative PCR (qPCR) reactions were conducted in triplicate, utilizing β-actin (Forward: TCCCTGGAGAAGAGCTACG, Reverse: GTAGTTTCGTGGATGCCACA) as the endogenous control. The comparative ΔCt (threshold cycle) method was employed to evaluate relative gene expression, and the primer sequences used for specific markers of odontoblast differentiation included NESTIN (Forward: GAAACAGCCATAGAGGGCAA, Reverse: TGGTTTTCCAGAGTCTTCAGTGA) and DSPP (Forward: TGACAGCAATGATGAGAGTG, Reverse: CACTGGTTGAGTGGTTACTG).

### Confirming Knockout at the Protein Level

Cells were lysed directly on the plate using a lysis buffer containing 20 mM Tris-HCl (pH 7.5), 150 mM NaCl, 15% glycerol, 1% Triton X-100, 1M β-glycerophosphate, 0.5M NaF, 0.1M sodium pyrophosphate, orthovanadate, PMSF, and 2% SDS. Following lysis, 25 U of Benzonase Nuclease (EMD Chemicals, Gibbstown, NJ) and a 100× phosphatase inhibitor cocktail were added to the lysate. To prepare the samples for analysis, 4× Laemmli sample buffer (Bio-Rad, #1610747), consisting of 950 μL of sample buffer and 50 μL β-mercaptoethanol (Sigma, #M7522), was added. The mixture was then heated at 95°C for 10 minutes. Subsequently, 15 μL of the protein sample was loaded onto an SDS-PAGE gel using a Protean TGX precast gradient gel (4%−20%) (Bio-Rad, #17000546) and transferred onto a nitrocellulose membrane (Bio-Rad, #1620115) using a semi-dry transfer system (Bio-Rad). The membranes were blocked for 1 hour with 5% BSA, followed by incubation overnight at 4°C with primary antibodies on a rocker. The primary antibodies used were AMBN (Santa Cruz #sc-271012, 1:500), SP6 (Atlas #HPA024516, 1:1000), DLX3 (Abnova #H00001747-M09, 1:500), DSPP (Santa Cruz #7363-2, 1:500), H3 (Abcam #Ab1791, 1:1000), and β-Actin (Cell Signaling #13E5, 1:10,000) all prepared in 5% BSA. The following day, the membranes were washed three times with 1× TBST at 10-minute intervals. Afterward, they were incubated for 1 hour at room temperature with an anti-rabbit IgG HRP-conjugated secondary antibody (Bio-Rad, #1721019) (1:10,000) and an anti-mouse IgG HRP-conjugated secondary antibody () prepared in 5% milk. After incubation, the membranes were rewashed with 1× TBST (three times, 10-minute intervals) and developed using the Immobilon Luminol reagent assay (EMP Millipore). Finally, protein bands were visualized using a Bio-Rad ChemiDoc Imager.

### Graphics and illustrations

The illustrations in the graphical abstract and in Figures were created with BioRender.com.

### Statistical analysis

All quantifications show the mean, and error bars are ± SEM. Ordinary one-way ANOVA was used for multiple comparisons. A two-tailed, unpaired t-test was used for comparing groups of two using GraphPad Prism. P-values < 0.05, 0.01, 0.001, 0.0001 are indicated with *, **, *** and ***, respectively. Software used and methods for analysis and quantification of each data in this manuscript are described in the method section.

---

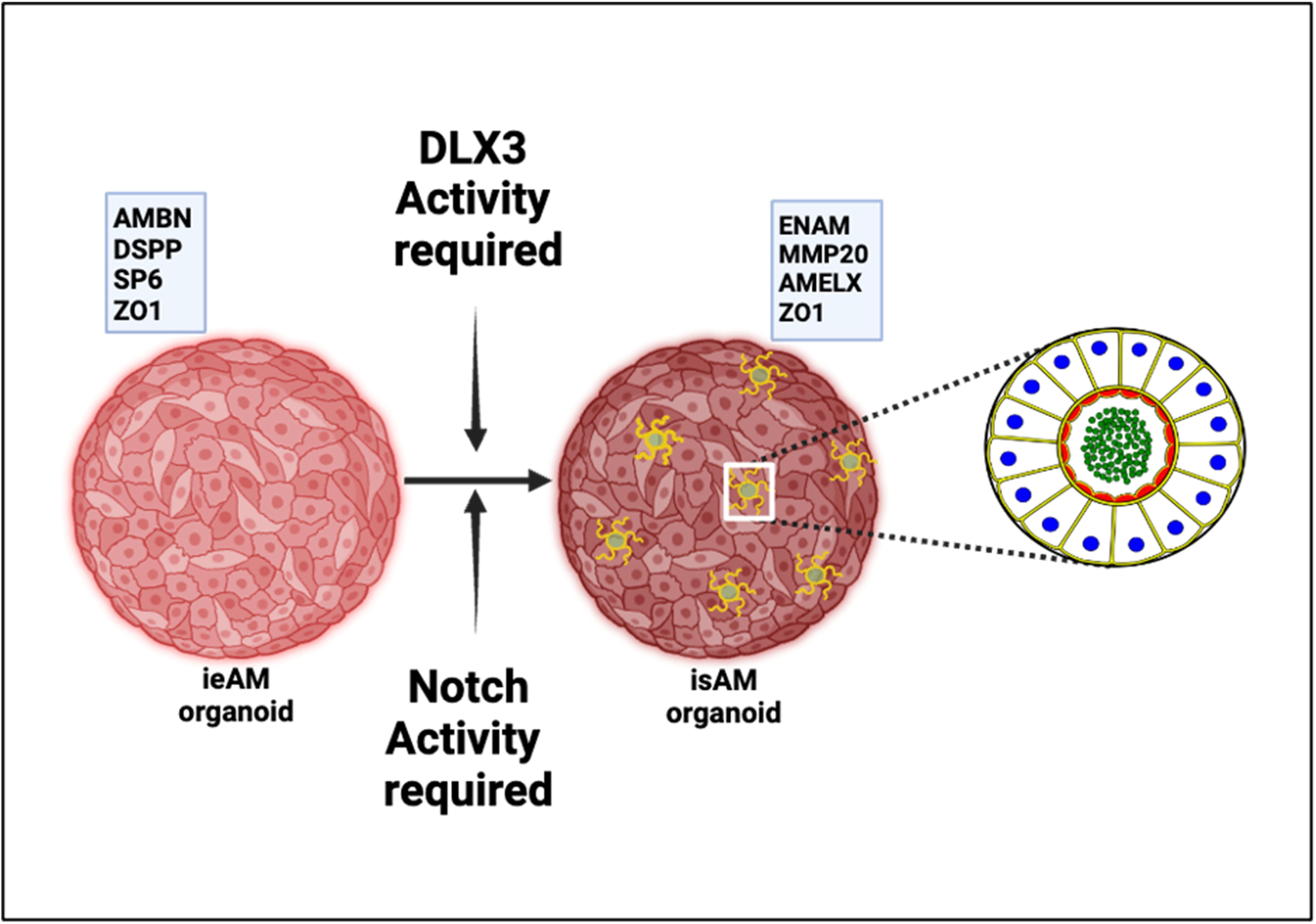

